# Mesocosm experiments reveal the loss of migratory tendencies in a recently isolated population of three-spined sticklebacks

**DOI:** 10.1101/2022.05.09.491155

**Authors:** A. Ramesh, J. Gismann, T.G.G. Groothuis, F.J. Weissing, M. Nicolaus

**Author notes:** Shared first authorship (A. Ramesh, J. Gismann).

## Abstract

In the 1970s, water management in the Netherlands resulted in numerous isolated populations of three-spined sticklebacks, which can no longer migrate from freshwater to the sea. We tested whether ∼50 years of isolation resulted in reduced migratory tendencies in these ‘resident’ sticklebacks. Lab-based individual testing showed behavioural divergence between residents and migrants, but also produced counter-intuitive results, especially with regards to movement tendencies. To detect differences in migration tendencies, we set up a semi-natural mesocosm, consisting of connected ponds, where movements of numerous individuals could continually be tracked at larger spatial scales. We found that wild-caught residents and migrants exhibited no differences in movement tendencies ‘within ponds’, but residents moved significantly less ‘between ponds’ than migrants. Between-pond movements were consistent and the observed differences were robust across contexts (changes in water flow and group size). Our study reveals that larger-scale movement tendencies can diverge over short time scales in response to human-induced isolation, and highlights the importance of observing behaviour in ecologically relevant setups that bridge the gap between lab and field studies.

## Introduction

Habitat fragmentation is one of the major threats for biodiversity, particularly for migratory species that depend on multiple habitats to complete their life cycle (1). In the north of the Netherlands, pumping stations have disrupted the connectivity between marine and riverine habitats, confining some fish populations to freshwater habitats without the possibility to migrate to the sea. Such forced isolation can cause rapid phenotypic responses and life-history changes (mammals and birds: (2); fish: (3–6)). Using individual lab-based assays, we have previously shown that this is indeed true for three-spined sticklebacks (*Gasterosteus aculeatus*): ‘resident’ populations, isolated for ∼50 years, were found to diverge in morphology and in behaviour from their ‘migrant’ ancestors (7), with part of the divergence having a genetic basis (8). Regarding movement-related behaviours, population differences uncovered in the lab were surprising at first because residents, that were expected to exhibit lower movement tendencies than migrants, were instead more active and more exploratory (7). We hypothesized at that time that this may be due to stress, induced by testing in social isolation, which might have affected wild-caught migrants disproportionately more than wild-caught residents, as migrants are thought to shoal extensively as an anti-predator strategy to higher predation risk in the open sea. Alternatively, small-scale experimental settings in the lab may not be suited to study larger-scale processes like migration. More generally, for wild-caught animals, lab conditions necessarily present a novel environment and fail to mimic natural complexity in biotic and abiotic factors, including the animals’ social environment (9–12). However, studying dispersal or migration behaviour in the field is often logistically challenging (especially in aquatic environments and for small fish) and frequently lacks data about the animals’ social groups (13).

To bridge the gap between lab and field studies, we set up a semi-natural mesocosm consisting of connected ponds, in which groups of fish can be remotely tracked over extensive periods of time. We here report the first experiment that aimed to test for consistent differences in movement tendencies between wild-caught ‘resident’ and ‘migrant’ sticklebacks and to disentangle the effects of spatial scale (within and between ponds), social environment (group size), and ecological conditions (water flow) on movement patterns. The results of the second experiment, aimed at disentangling genetic and non-genetic effects, are reported in (8). Under these experimental conditions, we tested (a) if residents and migrants exhibit differences in their movement tendencies, (b) if the spatial scale of movement matters, and (c) how consistent these patterns are under varying conditions (group size and water flow).

### Methods

### Mesocosm system

The mesocosm consists of two independent systems of five ponds (each Ø 1.6m, with water depth of 80cm), connected linearly with opaque corridors (each of length ∼1.5m and Ø 11cm), spanning a linear distance of ∼14m (Fig.1). The system is supplied with freshwater from a natural ditch, with the possibility of creating water flow (∼0.7cm/s), mimicking the wild conditions, which also acts a cue for migration (14). This system allowed to measure the movement of individual sticklebacks within and between ponds. The first pond (labelled 1 in Fig.1), enriched with plastic plants, was used to quantify within-pond movements, while the whole system of five connected ponds was used to record between-pond movement tendencies (see details in Supp. info.1).

**Figure 1:**
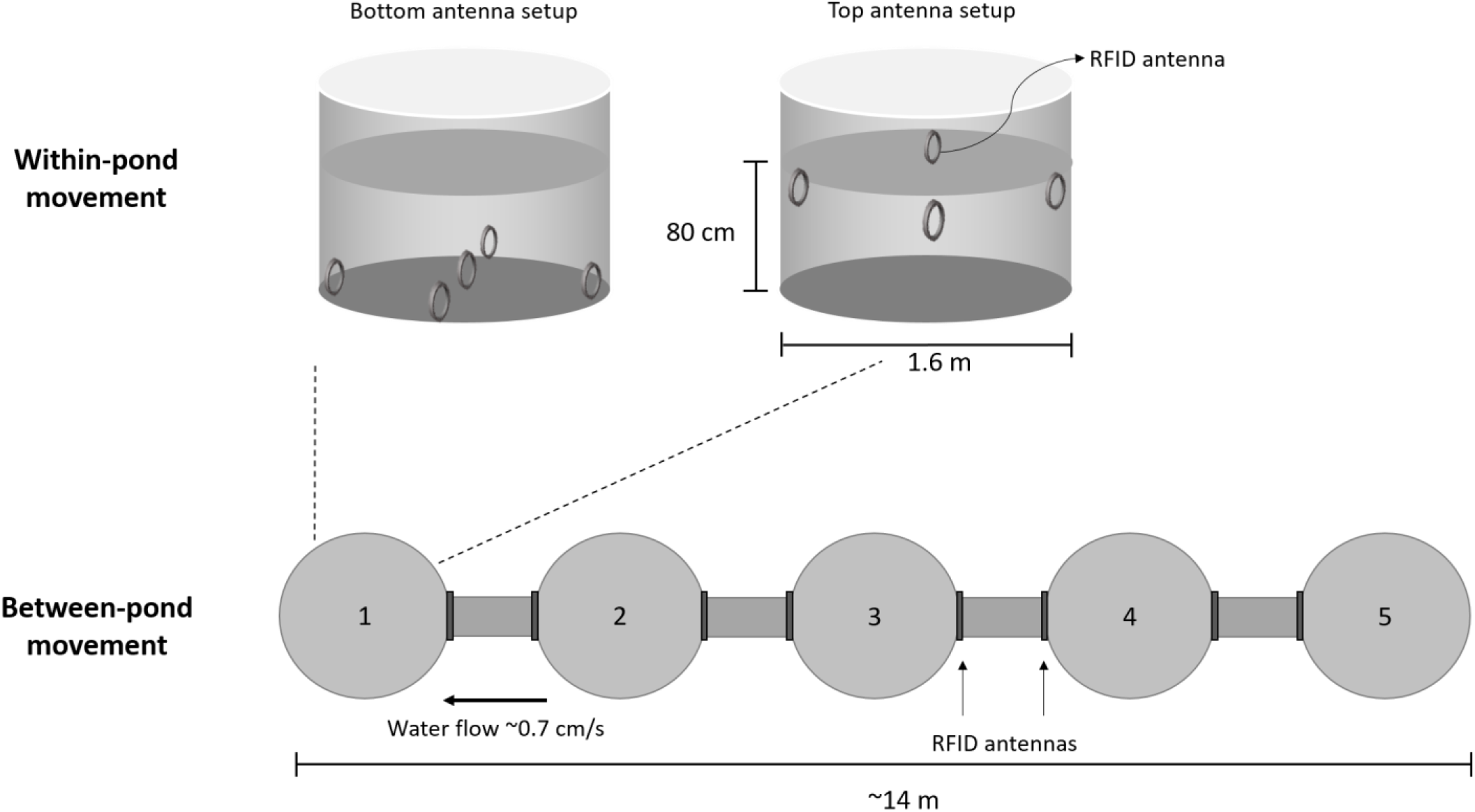
Experimental setup. The mesocosm consisted of two sets of five linearly connected ponds (1 to 5) equipped with circular RFID antennas that automatically detect crosses of PIT tagged individuals. Fish were released into pond 1. This pond was equipped with nine RFID antennas (five on the bottom and four on top of the water column), allowing us to quantify within-pond movements. The connections between adjacent ponds were equipped with two RFID antennas, allowing us to quantify the number and direction of movements between ponds.

We used a Radio-Frequency-Identification (RFID) system consisting of circular RFID antennas (Ø 10cm), data loggers and Passive Integrative Transponders (PIT tags) (*Trovan, Ltd*., *Santa Barbara, California*) to record movements of tagged sticklebacks (details in Supp. info.2). Nine circular antennas were placed in the first pond to record within-pond movements and two antennas were placed at both ends of each of the four connecting corridors to measure between-pond movement tendencies (Fig.1). Each antenna records the unique PIT-tag ID of the fish along with a time stamp, stored on a USB drive in the central data logger. The sensitivity of the system was set to three reads per second per unique tag. In a pilot study, we validated the reads using video recordings and found that it corresponded well with the entry and exit times of fish.

### Experiment-1

We created five groups of migrants and six groups of residents, each consisting of 10 randomly selected individuals (total: *Nmig*=49 and *Nres*=60). While we always tried to maintain the group size to 10 fish, tag-loss and other technical difficulties led to one group of migrants having nine fish and another with 11 fish. Groups were housed in separate small holding ponds for 24 hours before the start of the experiment. On the experimental day, one resident and one migrant group were released simultaneously (to avoid temperature or seasonal biases) into separate mesocosms. The individuals in each group were first monitored for within-pond movement by confining the fish to the starting pond for the first five hours (Fig.1) and then for between-pond movement for ∼16.5 hours, after opening the connection to the other ponds (Fig.1; Supp. info.2).

### Experiment-2

In a next step (after ∼one month), we combined all migrants and, separately, all residents (after excluding 12 fish which either had died or lost tags) into two large groups (*Nmig*=45, *Nres*=52) and quantified between-pond movements in these two groups in the same separate mesocosm setups over four days. In addition, we alternated flow and no-flow conditions on consecutive days (see Supp. info.1).

### Analyses

For each individual, we quantified within-pond movement as the number of times a fish crossed different bottom and surface antennas separately (Fig.1). We deemed the number of separate visits to a particular antenna unreliable for measuring movement patterns because fish that stayed longer near an antenna were recorded as multiple disconnected set of reads, as if they visited the antenna multiple times. Between-pond movement was quantified as the number of crosses a fish made through the corridors connecting two ponds (Fig.1). Fish that did not get detected by any antenna were given a score of zero crosses.

We then analysed if residents and migrants differed in the number of crosses for within-and between-pond movements (Experiment-1) and whether they were consistent across contexts (group size and flow) (Experiment-2). Briefly, we considered the number of crosses within or between ponds as response variable separately in univariate generalized linear mixed models with Poisson errors. In all models, we included *origin* (resident vs. migrant) as a fixed factor and *group-ID* and an observation-level *‘Obs’* (Observation-level random effects to control for overdispersion, (15)) as random effects. For Experiment-2, *treatment* (flow vs. no flow) and its interaction effect with *origin* were added as fixed effects and *individual-ID* as a random effect to account for individual repeats. Additionally, we analysed whether the fraction of fish that did not exit the first pond differed between migrants and residents using Fisher’s exact test. Repeatability and correlation of number of crosses across contexts were also calculated (Supp. info.3). All analyses were carried out in R v. 4.1.0, R Core Team (2019). For complete description of the analyses see Supp. info.3.

## Results

In Experiment-1, residents and migrants showed a broad distribution of number of crosses at both bottom and top antennas (Fig.2a, b) and the differences between the groups were in both cases not statistically significant (Table 1; Median bottom-antenna crosses: Residents=23, Migrants=14; Median top-antenna crosses: Residents=3.5, Migrants=8). In contrast, residents exhibited much lower numbers of crosses between ponds than migrants (Fig. 2c; significant effect of *Origin* in Table 1; Median pond crosses: Residents=0, Migrants=16). Furthermore, the proportion of ‘non-leavers’, i.e., individuals that did not exit the first pond, was significantly higher in residents than in migrants (55% in residents vs. 28.6% in migrants, odds ratio=3.02, p=0.007).

**Table 1.** Results of the statistical analysis of movement within and between ponds using generalised linear mixed models. Estimates of fixed effects (β) in log-scale are given with their 95% confidence intervals (CI) and variance components are given with their standard deviation. Fixed effects that significantly differ from zero are denoted in bold. Sample sizes experiment-1: *Nmig*=5 groups (49 individuals), *Nres*=6 groups (60 individuals); experiment-2: *Nmig*=1 group (45 individuals), *Nres*=1 group (52 individuals).

|  | Experiment-1 |  |  | Experiment-2 |
| --- | --- | --- | --- | --- |
|  | Bottom crosses | Top crosses | Pond crosses | Pond crosses |
| Fixed effects | $\beta$ (95% CI) | $\beta$ (95% CI) | $\beta$ (95% CI) | $\beta$ (95% CI) |
| Intercept | **2.61 (2.13, 3.08)** | **1.98 (0.30, 3.63)** | **1.90 (0.63, 3.13)** | **2.53 (1.87, 3.17)** |
| Origin<sup>1</sup> | 0.51 (-0.12, 1.15) | -0.68 (-3.03, 1.53) | **-2.26 (-4.04, -0.58)** | **-1.77 (-2.68, -0.87)** |
| Treatment<sup>2</sup> | - | - | - | -0.14(-0.44, 0.16) |
| Origin<sup>1</sup> × Treatment<sup>2</sup> | - | - | - | **-0.72 (-1.18, -0.27)** |
| Random effects | Var (sd) | Var (sd) | Var (sd) | Var (sd) |
| Group-ID | 0.11 (0.33) | 2.94 (1.72) | 0.95 (0.98) | - |
| Obs | 1.21 (1.10) | 1.14 (1.07) | 5.02 (2.24) | 0.81 (0.90) |
| Individual-ID | - | - | - | 4.11 (2.02) |

<sup>1</sup>'migrant' is used as reference category; <sup>2</sup>'flow' is used as reference category

**Figure 2:**
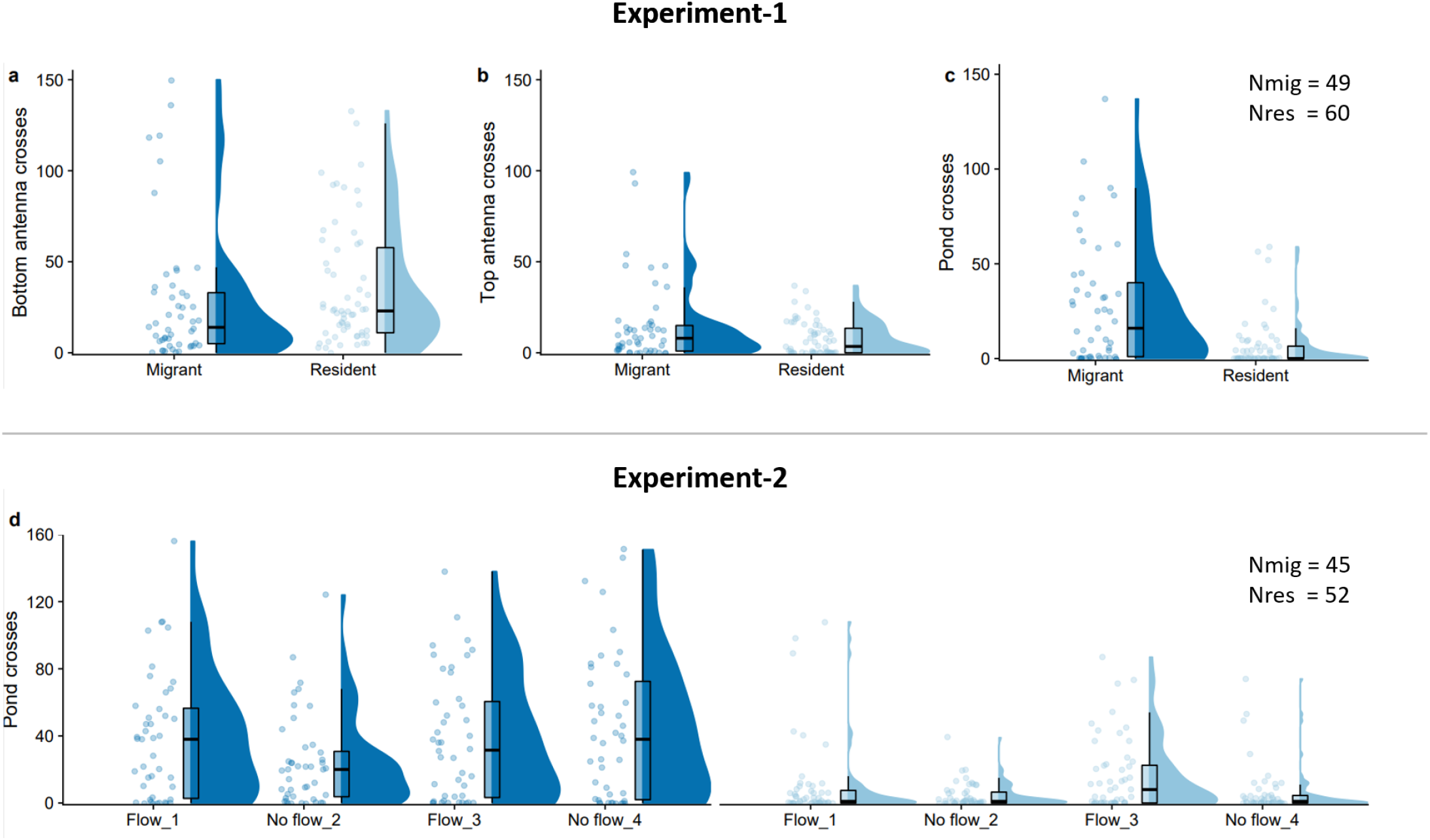
Within-pond and between-pond movement of resident and migrant sticklebacks. a-b) within-ponds crosses at the bottom and top antennas respectively (Experiment-1); c) between-pond crosses in Experiment-1; d) between-pond crosses in relation to the daily flow treatment in Experiment-2. In all graphs, individual crosses (dots), boxplots and density kernels are shown for migrant (dark blue) and resident (light blue) sticklebacks.

In Experiment-2, as in Experiment-1, residents moved consistently less between ponds than migrants (Fig.2d). Furthermore, fish moved more between ponds in the presence of flow and the trend was slightly stronger for residents than migrants (Fig.2d; significant *Origin*×*Treatment* effect in Table 1). Individual movement tendency between ponds was moderately repeatable across ecological contexts but very weakly correlated over social contexts (Supp. info. 3.). However, we clearly see from Figure 2 and Table 1 that the difference between residents and migrants was maintained across different contexts.

## Discussion

We have previously shown that ∼50 years of isolation potentially led to rapid behavioural and morphological divergence of residents from migrants (7), which mimics the divergence observed in another long-isolated population of sticklebacks (16). Both studies assayed individual movement tendencies under artificial housing conditions in the lab and showed counter-intuitive patterns: residents showed either higher (7) or inconsistent patterns (16) in activity/exploration levels compared to migrants. Here, we show that the same populations as in (7) exhibited movement tendencies as predicted previously, when they were tested in a semi-natural setting (relevant social/ecological context and spatial scale): Resident populations exhibited lower movement tendencies than their migrant counterparts. These differences, detected only at large spatial scale, remained consistent across ecological and social contexts. Together with the previous results on F1 lab-born juveniles (8), this study suggests that our mesocosm setup, by allowing water flow, testing in groups and larger spatial scale (∼15m length), is much better suited to characterize individual movement patterns related to migratory behaviour than lab-based assays in social isolation in small tanks.

Our study reveals that the detection of population differences in stickleback behaviour was scale-dependent (only detectable between, but not within ponds). This is probably because in the wild, sticklebacks exhibit considerable foraging movements over days (median of 40m upstream, (17)) and hence their within-pond movements, representing foraging movements, may not differ between populations. However, wild migrants in our field system travel 10s of kilometres inland within a few days (Pers. comm. from water authorities) and thus require sufficient space and navigation cues (e.g. flow velocity (14)) to express their natural behaviour.

Tests in the lab, though invaluable for studies on animal behaviour owing to controlled settings, are not without drawbacks. Firstly, they cannot offer the more natural conditions mentioned above (e.g. spatial scale, appropriate social or ecological contexts), which may be particularly important for wild-caught animals. They may constrain the level of behavioural expression to some extent, such as the ‘freezing’ behaviour of wild-caught migrants in our previous studies (7). Reassuringly, we observed that this was much less of an issue for lab-bred animals: lab-born F1 juveniles did not freeze in lab tests and their movement-related behaviours measured in the lab and in the mesocosms positively correlated (Fig.S1). Secondly, lab-tests are performed in highly-controlled or novel setups. This can lead to homogenization of behavioural expression (e.g. decreased variance over time (18)) or uncovering ‘cryptic’ behavioural variation (with novel behaviours and increased variance in behavioural expression (19)). We thus advocate using mesocosms or other semi-natural setups (e.g. 20–27), to bridge lab and field studies. They circumvent the mentioned drawbacks and provide valuable insights undetectable in classical behavioural setups, especially for wild populations.

Our results further support the idea that forced isolation in freshwater is followed by phenotypic changes as reported for sticklebacks isolated after the last glacial retreat (e.g. reduction in lateral plates and reduced swimming abilities (28–30)). Many of these morphological and behavioural changes are underlined by genetic differentiation and are true adaptations to a resident lifestyle (31,32). Additionally, we show that freshwater-induced phenotypic changes in sticklebacks can occur even on contemporary timescales (see also (33–35)) and can have a genetic component (8). Residents in our study populations are thus likely on a trajectory to losing their migration tendencies and already (partially) adapted to complete residency. Current conservation management includes building fishways to reconnect land-locked and migratory populations. In this context, it is important to consider that residents may be less likely to use fishways due to lowered migration tendencies. This may require a revision in the evaluation criteria for the success of these conservation efforts. An exciting future avenue will be to study to what extent and how quickly individual migration tendencies will be affected when the two populations reconnect.

## Ethics

Wild animals were sampled using a fishing permit from *Rijksdienst voor Ondernemend Nederland* (the Netherlands) and an angling permit from the Hengelsportfederatie Groningen-Drenthe. Housing and testing of behaviours were in adherence to the project permit from the *Central Committee on Animal Experiments* (CCD, the Netherlands) under the licence number *AVD1050020174084*.

## Data accessibility

Datasets are provided as electronic supplementary methods and we will be upload it to a data repository after acceptance.

## Authors’contributions

AR: conceptualization, data collection, data curation, analysis, writing an original draft of the manuscript, review and editing the manuscript; JG: conceptualization, data collection, data curation, analysis, writing: review and editing the manuscript; MN: conceptualization, analysis, supervision and writing: review and editing the manuscript; FJW & TGGG: conceptualization, supervision and writing: review and editing the manuscript. All authors declare no competing interests and gave final approval for publication and agreed to be held accountable for the work performed herein.

## Acknowledgments

We thank Dennis de Worst and Willem Diderich for help with fish care and advice on experimental design and other animal caretakers for looking after the sticklebacks. We thank Peter Paul Schollema from the Water Authorities Hunze en Aa’s and Jeroen Huisman from van Hall Larenstein, University of Applied Sciences for help with catching sticklebacks.

