## Supplementary information for "Mesocosm experiments reveal the loss of migratory tendencies in a recently isolated population of three-spined sticklebacks"

**Supplementary information 1: Description of the mesocosm and the tracking setup**

*Experiment-1*

*Within-pond movements*

On the morning of testing (~10 a.m.), one test group of each origin was released into the first pond that was temporary disconnected from the other ponds by a cap blocking the entrance to the corridor. There was no water flow when recording within-pond movement tendencies. Five circular antennas were placed upright on the bottom of the pond (“bottom antennas”), and four antennas were placed just below the water surface (“surface antennas”) (Fig. 1). To assess within-pond movements, we computed crosses that an individual made between bottom antennas or the surface antennas separately. Crosses that were made between a bottom and surface antenna were excluded as these hardly occurred. The experiment lasted for five hours.

*Between-pond movements*

After five hours, we gently removed all antennas from the first pond. At this point we also turned on the flow in the system to create a cue for migration (Fig. 1c). Fish were given 30 minutes to recover from the disturbance caused by removing the antennas after which the connection from pond 1 to the other ponds was gently opened. We then recorded the movement of fish between the five connected ponds (“crosses”) for the next 16.5h (~3.30 p.m. -8 a.m.). At the end of the experiment, fish were returned to their original smaller housing ponds. Testing all 5 migrant and 6 resident groups took place over a week (temperature ranged between 12^0^C and 15^0^C). All fish were checked at the end of the experiment to see if they still carried the tags and if the tags functioned correctly.

*Experiment-2*

*Between-pond movements*

Two weeks after we finished recording each individual for movement tendencies as above, we created one large group each of migrant and resident by combining all the fish (*Nmig* = 1 groups, 45 individuals; *Nres* = 1 group, 52 individuals) and monitored only the movement tendencies between-ponds simultaneously for the two groups and continuously for four consecutive days. During the study period, we furthermore alternated days with and without water flow (flow turned on / off at 10:00 AM each day and hence kept in that condition for ~24 hours). The flow treatment allowed testing whether the populations react differently to the presence of a migration cue.

**Supplementary information 2: Study populations and housing of fish**

We caught incoming migrants at a sea lock at the mouth of a river in Nieuwe Statenzijl (“NSTZ”; 53013’54.49’’, 7012’30.99’’), and resident sticklebacks in an adjacent land-locked polder (“LL-A”; 53017’56.14’’, 702’1.28’’) in the province of Groningen, The Netherlands (1). Fish were caught at the onset of inland migration, over a period of four weeks in March and April 2020. Fish of ≥4 cm in total length (from the tip of the snout to the tip of the tail) were transported to the lab in aerated plastic bags within two hours of capture. After acclimatization, fish were housed in groups of 25, separated by their origin (migrant or resident), for a week prior to experimentation in small holding ponds (~ 100 L tanks filled with freshwater from a nearby ditch) under natural temperature and light conditions. Fish were fed a mixture of brine shrimps and bloodworms (3F Frozen Fish Food bv.), once a day, ad libitum. Fish were tagged with 8 mm Passive Integrated Transponders (PIT tag; Trovan, Ltd., Santa Barbara, California) for individual identification, under anaesthetization in buffered MS-222 solution (0.25 – 0.3 g/L, pH=7.5-8.0). PIT tags were injected in the abdominal cavity (following (2)). Before experiments, all fish were allowed at least five days of recovery in the housing pond with the same group. Mortality rate after PIT tagging was very low (<1% in the first week).

**Supplementary information 3: Estimating consistency of between-pond movements.**

To quantify individual consistency in between-pond movements across ecological contexts (flow/no flow), we ran univariate generalised linear mixed model (GLMMs) with Poisson errors using the dataset from Experiment-2 and the lme4 package (3). For repeatability across social environments (small vs large group size), we combined the crosses data from Experiment-1 and day 1 and 3 of Experiment-2 with flow.

We used the number of crosses between ponds as the response variable, with *origin* (resident vs. migrant), *treatment* (social context: small vs large group size or ecological context: flow vs no-flow in two separate models) and their interaction (*origin* × *treatment*) as a fixed factors and *individual-ID* as a random effect. In addition, we added *Obs* as observation-level random effects to control for over-dispersion (OLRE, (4)). We used these ‘full’ and ‘simplified’ models (omitting all the fixed effects) to calculate ‘adjusted’ and ‘raw’ repeatabilities respectively. Repeatabilities are defined as the the ratio of among-individual variance (Vind) to total variance (Vtotal = Vind + Vresidual). We calculated repeatabilities in their original scale, along with their confidence intervals using the ‘rpt’ function with 1000 bootstraps using the ‘rptR’ package (5). We were not able to calculate repeatabilities for different social contexts due to lack of model convergence. Hence we resorted to using Spearman correlation as the data is not normally distributed. All analyses were carried out in R v. 4.1.0, R Core Team (2019).

Between-pond movement was moderately repeatable across ecological contexts (Adjusted R(95% CI) = 0.42 (0.34, 0.51) and Raw R(95% CI) = 0.38 (0.30, 048)). Across social context, individuals were not very consistent with low correlation coefficients (Spearman ρ = 0.35, p < 0.001). This could be because timescale and sample size were not balanced between Experiment-1 and -2. While repeated data were collected over consecutive days in Experiment-2, single data points were collected a month apart in Experiment-1. However we see that the residents were consistently moving less than migrants in all contexts.

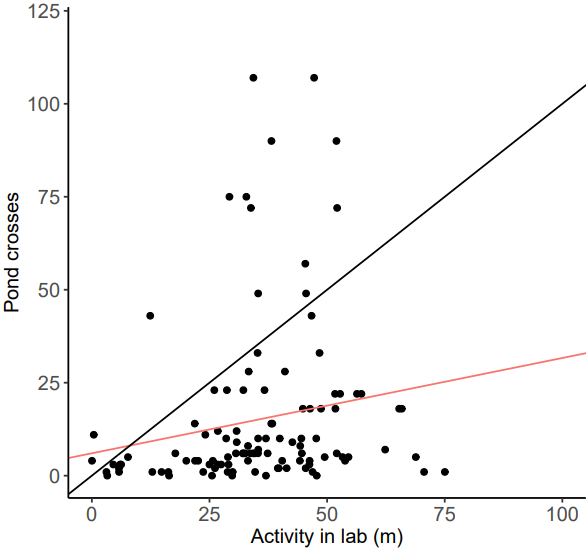

**Figure S1: Correlation of movement tendencies of lab-raised F1 migrants, residents and hybrids tested in the lab and in the mesocosm.** In a separate experiment and set of animals, (F1 sticklebacks raised in the lab from (1)), we performed both, an activity assay in the lab, where individual fish were assessed for general movement tendencies for 20 minutes in their home tank (30 x 16 x 18 cm (L x W x H)) (according to methods in (6)) and movement tendencies across-ponds in the mesocosm as in Experiment-1. Black line represents the identity lines, x = y. The red line is the ordinary least squares regression line. Lab-based activity (total distance covered in meters in 20 mins) and number of pond crosses in the mesocosm were positively and significantly correlated (Spearman ρ = 0.33, p < 0.01).
